# Quercetin as a Bitter Taste Receptor Agonist with Anticancer Effects in Head and Neck Cancer Cells

**DOI:** 10.1101/2025.09.17.676947

**Authors:** Gavin Turner, Sarah M. Sywanycz, Brianna L. Hill, Robert D. Wardlow, Robert J. Lee, Ryan M. Carey

## Abstract

**Background/Objectives:** Quercetin is a bitter compound with demonstrated anticancer effects in preclinical models of head and neck squamous cell carcinoma (HNSCC). In taste transduction, bitter compounds activate bitter taste receptors (T2Rs), a group of G protein-coupled receptors with downstream signaling that includes cytosolic calcium (Ca^2+^) release. T2Rs are expressed in HNSCC cells, where their activation induces apoptosis *in vitro*. Increased T2R expression in HNSCC also correlates with improved patient survival. The objective of this study was to investigate the role of quercetin as an anticancer T2R agonist in HNSCC cells *in vitro* and *ex vivo*.

**Methods:** Quercetin-mediated Ca^2+^ responses were assessed using live cell Ca^2+^ imaging in the presence of the T2R14 antagonist LF1 and G-protein inhibitor YM-254980 (YM) in UM-SCC-47 and FaDu HNSCC cell lines. Cell viability was evaluated using crystal violet assays in cell lines and MTS assays in patient-derived tumor slices. Mitochondrial depolarization was measured with TMRE in the presence and absence of T2R pathway inhibitors.

**Results:** Quercetin induced a Ca^2+^ response in HNSCC cells, which was significantly reduced by LF1 and YM. Quercetin also decreased cell viability *in vitro* and showed a potential decrease in viability in tumor slices but was not statistically significant. Finally, quercetin caused mitochondrial depolarization, which was reduced in the presence of LF1 but not by YM.

**Conclusions:** In HNSCC cells, quercetin causes a Ca^2+^ response that is likely mediated by T2R14, decreases viability, and causes mitochondrial depolarization.

## 1. Introduction

Head and neck cancers of the upper aerodigestive tract had a combined global incidence of over 890,000 cases in 2022 with projections rising to 1.37 million cases by 2045 [1, 2]. Natural compounds are being studied as potential anticancer agents, with plant flavonoids showing promise in pre-clinical models [3, 4]. Among them, quercetin – a flavanol widely distributed in fruits, vegetables, and medicinal plants – has been extensively investigated [4–6].

Quercetin is a bitter compound [7]; bitter compounds bind to bitter taste receptors (T2Rs), a family of G protein-coupled receptors (GPCRs) with approximately 25 isoforms in humans [8–10]. Canonical T2R signaling involves decreases in cAMP, increases in IP_3_, release of endoplasmic reticulum calcium (Ca^2+^) stores, and opening of TRP channels, leading to taste cell depolarization [11–20]. Beyond taste transduction, T2Rs are expressed extra-orally and have various actions: modulating innate immunity in airway epithelial cells via nitrous oxide production and increased ciliary beating, effecting hormone signaling in the gut through GLP-1 secretion, and promoting bronchodilation in airway smooth muscles via large-conductance Ca^2+^-activated K^+^ channel activation [21–24]. Importantly, T2Rs are also expressed in solid tumors, including HNSCC, where their activation has been linked to proteosome inhibition, apoptosis, and possibly mitochondrial Ca^2+^ overload and depolarization [25–27]. Increased T2R expression in HNSCC tumors correlates with improved patient survival, supporting a potential role for T2Rs as therapeutic targets [25]. Notably, the T2R14 agonist lidocaine has demonstrated anticancer effects [28–30], leading to clinical trials evaluating its efficacy for solid tumors such as breast cancer [31] and HNSCC [32].

Quercetin was initially shown to activate T2R14 and cause intracellular Ca^2+^ responses [33]. Subsequent studies confirmed quercetin binding to multiple T2Rs in HEK 293T cells and other cell types, including human choroid plexus papilloma cells and enteroendocrine L cells [34–36]. While these findings support quercetin’s role as a bitter agonist, further exploration is warranted to clarify its functional effects, signaling mechanisms, and target cell types. In this study, we show that quercetin induces intracellular Ca^2+^ responses in HNSCC cell lines that are likely T2R-mediated. We build upon prior evidence of quercetin’s anticancer activity, demonstrating its ability to decrease HNSCC cell viability *in vivo* while exploring its effects in *ex vivo* tumor slices. Finally, we explore mitochondrial dynamics to evaluate the relationship between T2R activation and mitochondrial depolarization.

## 2. Materials and Methods

### 2.1 Cell Lines and Cell Culture

Human HNSCC cell lines included UM-SCC-47 ([SCC47] MilliporeSigma, St. Louis, MO USA; RRID: CVCL 7759) and FaDu (ATCC, Manassas, VA, USA; RRID: CVCL 1218). SCC47 is an HPV-positive HNSCC cell line derived from a tongue tumor of a 53-year-old male [37, 38]. The FaDu cancer cell line was derived from an HPV-negative tumor of the hypopharynx in a 56-year-old male [39]. Cells were cultured in high glucose Dulbecco’s Modified Eagle’s Medium (DMEM; Corning, Glendale, AZ, USA) supplemented with 10% FBS (Genesee Scientific, El Cajon, CA, USA), 1% penicillin/streptomycin (Gibco, Gaithersburg, MD, USA), and 1% nonessential amino acids (Gibco) at 37°C with CO_2_ 5%.

### 2.2 *Ex vivo* Tumor slices

Tumor tissue was acquired from HNSCC patients undergoing oncologic surgeries after written informed consent and approval of the University of Pennsylvania Institutional Review Board (IRB #417200). Specimens were embedded in agarose and sectioned into 300 µM slices using a vibratome (Precisionary Compresstome Vibratome VF-510-0Z; oscillations 6 and speed 3). Slices were cultured free-floating in high glucose DMEM supplemented with 10% FBS, 1% penicillin/streptomycin, and 1% nonessential amino acids at 37 °C with 5% CO_2_.

### 2.3 Reagents

Quercetin (Caymen Chemical Company, Ann Arbor, MI USA) 200 mM stock solutions were prepared in DMSO, aliquoted, and stored at −20 °C. Stocks were diluted into room temperature Hank’s Balanced Salt Solution (HBSS) for live cell imaging or into 5% CO_2_-equilibrated high glucose DMEM (with 10% FBS, 1% penicillin/streptomycin, and 1% nonessential amino acids) warmed to 37°C for viability assays. A maximally saturated working solution of quercetin was obtained by dissolving quercetin in HBSS at ≤500 µM with vortexing and shaking, followed by centrifugation to remove any small precipitates.

The T2R14 antagonist LF1 (Enamine, Monmouth Jct., NJ USA; Cat. No. Z85879385) was prepared as a 500 mM stock in DMSO, stored at −20 °C, and diluted to 500 µM in HBSS for experiments [40]. The Gα_q/11_ inhibitor YM-254980 ([YM] Caymen Chemical Company, Ann Arbor, MI USA) was prepared as a 10 mM stock in DMSO, stored at −20 °C, and diluted to 1 µM in HBSS for experiments.

### 2.4 Live Cell Imaging

Live cell imaging was performed using an Olympus IX-83 microscope (20x 0.75 NA PlanApo objective) with appropriate filters (Semrock, Rochester, NY), an Orca Flash 4.0 sCMOS camera (Hamamatsu, Tokyo, Japan), MetaMorph Microscopy Automation and Image Analysis Software version 7.10.1.161 (Molecular Devices, San Jose, CA USA), and XCite 120 LED Boost (Excelitas Technologies, Pittsburgh, PA). HBSS was used for fluorophore loading, washes, imaging buffer, and experimental/control conditions. HBSS contained LF1 or YM for experiments involving antagonists/inhibitors. Separate wells per plate were treated as technical replicates. Repeating experiments with different cell passages were treated as biologic replicates.

#### 2.4.1 Intracellular Ca^2+^ Dynamics

Cells were plated on 8-well glass bottom chamber slides (Cellvis, Mountain View, CA USA). The next day, cells were incubated in Calbryte 590 acetoxymethyl ester (AM) (AAT Bioquest, Pleasanton, CA USA) in HBSS for 45-60 minutes in the dark at room temperature, then washed and maintained in imaging buffer as described in section 2.4 Live Cell Imaging. TRITC filters (554/23 nm_ex_, 609/54 nm_em_, 573 nm long-pass dichroic; Semrock LED-TRITC-A-OMF) were used. Imaging was performed with a 20x (0.8 NA) objective with 150 ms exposure of and 25% LED intensity at 15 second intervals. Baseline recordings were obtained before treatments.

#### 2.4.2 Detecting autofluorescence

Cells were plated on 8-well glass bottom chamber slides. The next day, cells were washed with HBSS and maintained in HBSS for imaging. FITC (474/27 nm_ex_, 525/45 nm_em_, 495 nm long-pass dichroic; Semrock LED-FITC-A-OMF) and TRITC filters (described above) were used. Imaging was performed with a 20x (0.8 NA) lens, 25% LED intensity, and 150 ms exposure at 45 second intervals for 22.5 minutes. Baseline images were acquired prior to treatments.

#### 2.4.3 Mitochondrial Membrane Potential

Cells were plated on 8-well glass bottom chamber slides. The next day, cells were incubated in HBSS with or without antagonist/inhibitor for 45 minutes, followed by incubation with tetramethylrhodamine ethyl ester (TMRE) and Hoechst Dye (Caymen Chemical Company, Ann Arbor, MI USA) for 15 minutes in the dark at room temperature. Cells were washed and maintained in imaging buffer as described in section 2.4 Live Cell Imaging. DAPI filters were used to initially identify nuclei, and TRITC filters were used for imaging. Cells were imaged at 20x with 50 ms exposure and 25% LED intensity at 20 second intervals.

### 2.5 Viability Assays

Cell viability of cell lines was assessed using crystal violet. Cells were seeded onto 24-well plastic-bottom plates (Fisher Scientific) at 15% confluency for 48-hour experiments and 30% for 24-hour experiments and incubated at 37 °C with CO_2_ 5%. The following day, media was aspirated and replaced with quercetin-containing media or control media. After 24 or 48 hours, cells were washed with PBS, stained with crystal violet (0.1% in deionized water with 10% acetic acid), air-dried, dissolved in 30% acetic acid, and absorbance measured at 590 nm using a Tecan Spark 10M (Mannedorf, Switzerland). Technical triplicates were performed per condition with at least three biological replicates.

Cell viability of tumor slices was assessed using the 3-(4,5-dimethylthiazol-2-yl)-5-(3-carboxymethoxyphenyl)-2-(4-sulfophenyl)-2H-tetrazolium (MTS) assay. Slices were transferred to individual wells of a 96-well plate (Fisher Scientific) containing 100 µL of phenol-free, high glucose DMEM (supplemented with 10% FBS, 1% nonessential amino acids, and 1% penicillin/streptomycin) with or without 200 µM quercetin, followed by incubation for 24 hours. For each patient, three slices per condition were used, with three wells per condition lacking slices to serve as blanks for background subtraction. At 22 hours, CellTiter 96 Aqueous One solution MTS Cell Viability dye (Promega, Madison, WI, USA) was added to each well. At 24 hours, supernatant was transferred to a new plate and absorbance analyzed at 490 nm using a Tecan Spark 10M.

### 2.6 Statistical Analysis

All analyses were performed using GraphPad Prism version 10.3.1 (GraphPad Software, Boston, Massachusetts). When comparing two variables, unpaired t test was performed. Welch’s correction was performed when F test showed that the standard deviations of groups being compared were significantly different. When comparing multiple variables, ordinary one-way ANOVA followed by Tukey’s multiple comparisons test comparing the mean of each column to the mean of every other column was performed. Brown-Forsythe and Welch ANOVA test was performed when Brown-Forsythe Test showed that the standard deviations of groups being compared were significantly different. This was followed by Dunnet’s T3 multiple comparisons test comparing the mean of each column to the mean of every other column. For comparing viability of tumor slices, Wilcoxon matched-pairs signed rank test was used. Statistical significance for all experiments was set at P < 0.05.

## 3. Results

### 3.1. Quercetin Causes an Intracellular Ca^2+^ Response that is Likely T2R-mediated

Live-cell imaging with Calbryte 590 AM was used to measure cytosolic Ca^2+^ dynamics in response to quercetin in SCC47 and FaDu cells. Maximum normalized fluorescence signals (F/F_0_) were significantly greater in quercetin-treated conditions compared to control (HBSS vehicle only) in both cell lines, with average traces illustrating differences in signals over time (**Figure 1**). These findings indicate that quercetin induces an intracellular Ca^2+^ response in HNSCC cells. To determine whether these Ca^2+^ responses were T2R-mediated, experiments were repeated in the presence of T2R14 antagonist LF1 [26, 40]. Peak F/F_0_ signals after quercetin exposure were significantly reduced in LF1-treated cells compared to control in both cell lines (**Figure 1**), suggesting that T2R14 contributes to these Ca^2+^ responses.

**Figure 1.**
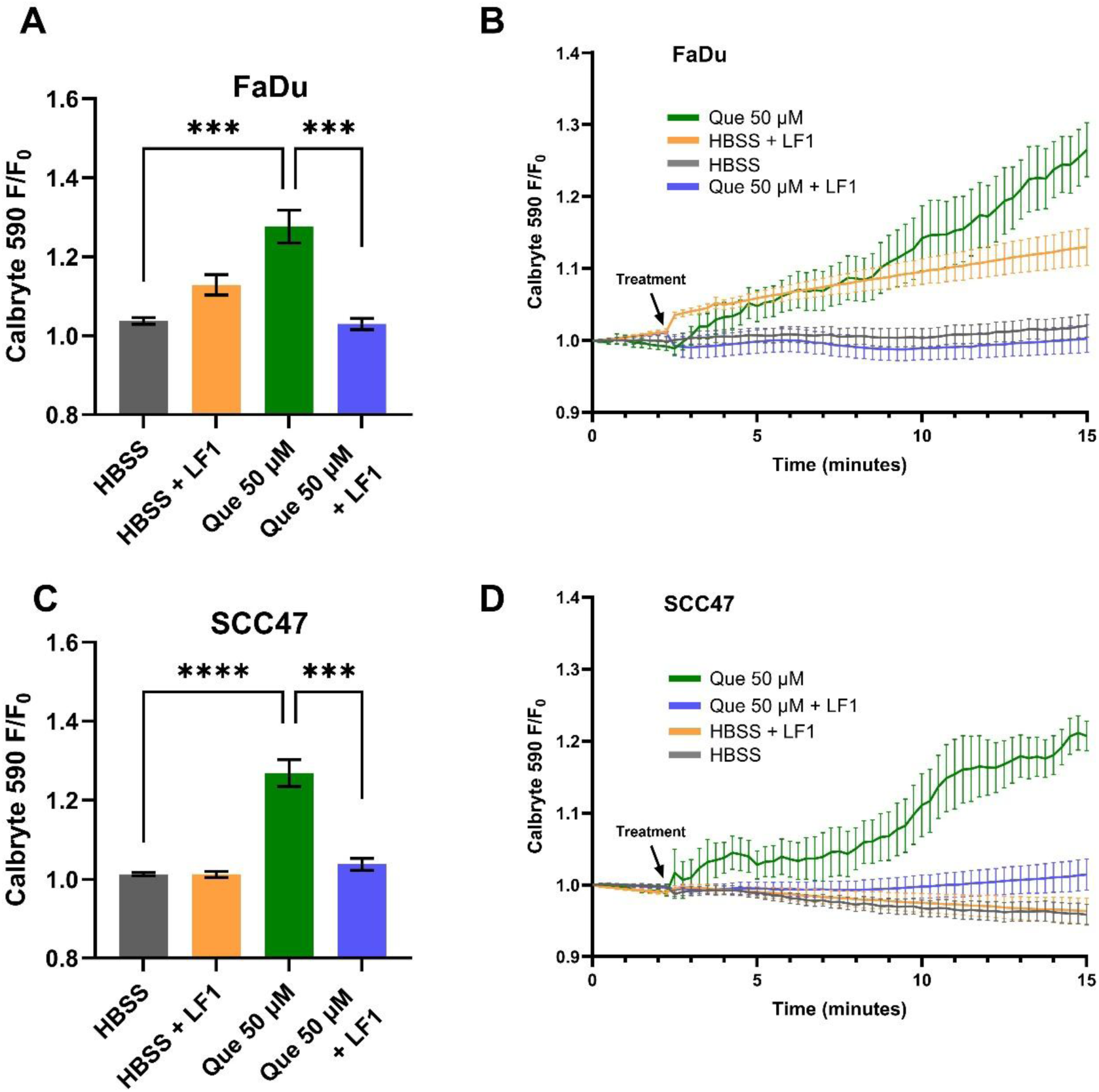
HNSCC cell line Ca^2+^ responses with quercetin in the presence and absence of T2R14 antagonist. Cells were incubated in Calbryte 590 AM with and without T2R14 antagonist LF1 prior to imaging. Quercetin with and without LF1 were added after baseline images were obtained; either HBSS or HBSS with LF1 served as the negative control. Cells were imaged at 20x using TRITC filters at 15 second intervals. Increased Calbryte 590 signal corresponds to increased intracellular Ca^2+^. (**A**) Peak F/F_0_ and (**B**) average traces with and without LF1 in FaDu cells. (**C**) Peak F/F_0_ and (**D**) average traces with and without LF1 in SCC47 cells. Experiments were conducted with technical replicates with n = 4 and repeated with biologic replicates n = 3. The statistical method used was Brown-Forsythe and Welch ANOVA with Dunnet’s T3 multiple comparisons test. Data points and error bars represent mean ± SEM. ***P < 0.001; ****P < 0.0001.

To further test if the Ca^2+^ response is GPCR-mediated, live-cell imaging was conducted in the presence of YM, a Gα_q/11_ inhibitor [41]. In SCC47 cells, maximum normalized fluorescence signals (F/F_0_) were significantly lower with YM treatment compared to control (**Figure 2**). A similar effect was observed in FaDu cells, where endpoint fluorescence signals after approximately 11 minutes of stimulation were signficantly reduced in YM-treated cells relative to control (**Figure S1**).

**Figure 2.**
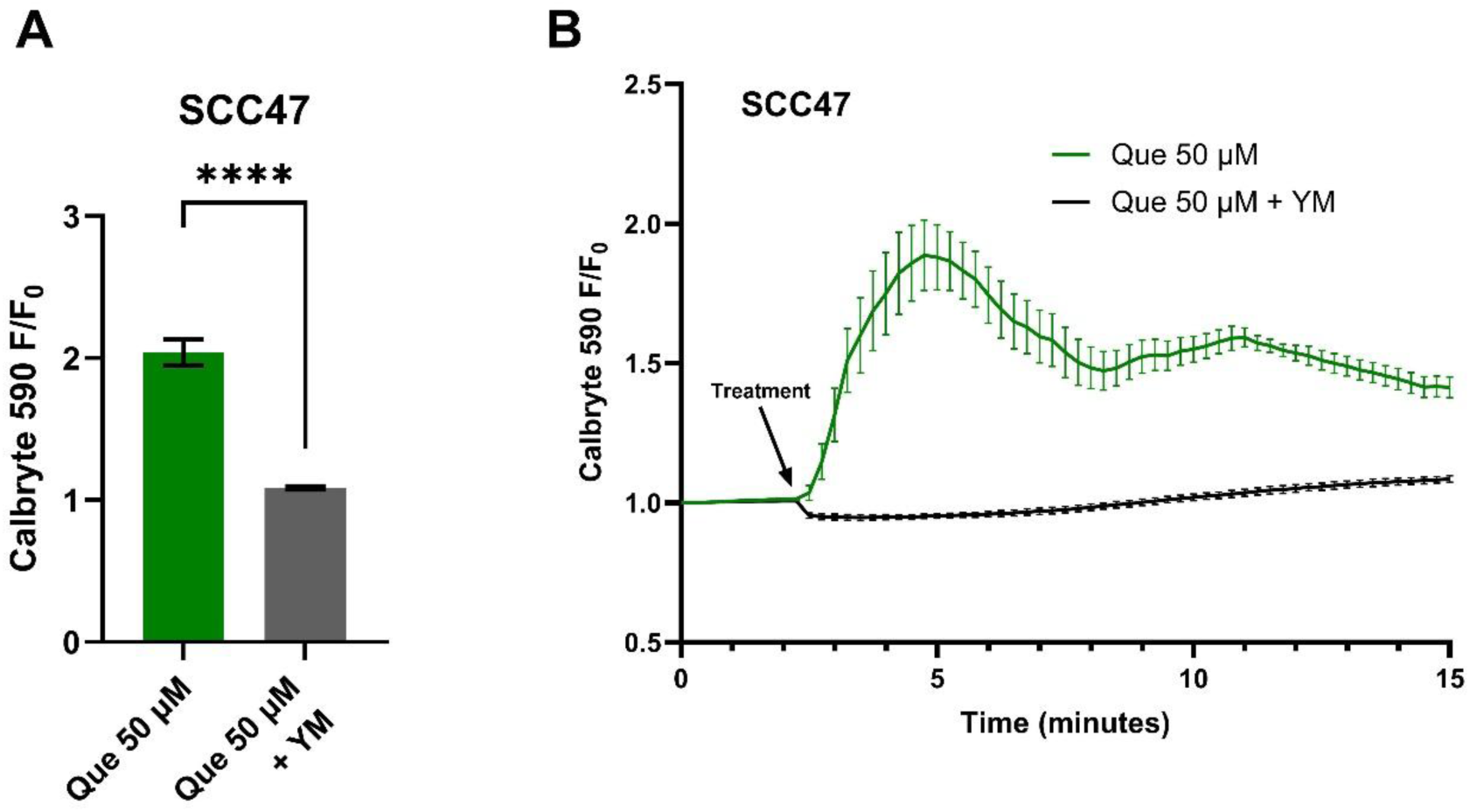
HNSCC cell line Ca^2+^ responses with quercetin in the presence and absence of Gα_q/11_ inhibitor. SCC47 cells were incubated in Calbryte 590 AM with and without YM prior to imaging. Quercetin with and without YM were added after baseline images were obtained; HBSS with or without YM served as the negative controls respectively. Cells were imaged at 20x using TRITC filters at 15 second intervals. Increased Calbryte 590 signals correspond to increased intracellular Ca^2+^. (**A**) Peak F/F_0_ with and without YM and (**B**) average traces in SCC47 cells are shown. Experiments were conducted with technical replicates with n = 4 and repeated with biologic replicates n = 3. The statistical method used was Welch’s t test. Data points and error bars represent mean ± SEM. ****P < 0.0001.

### 3.2. Intracellular Autofluorescence after Addition of Quercetin

To investigate the autofluorescent properties of quercetin in HNSCC cells, SCC47 and FaDu cells in the absence of fluorescent indicators were exposed to quercetin and imaged using FITC and TRITC filters. Upon addition of quercetin when using FITC filters, an immediate increase in fluorescent signals was observed and sustained over the approximate 23 minutes of imaging in both cell lines; no increases in fluorescence were observed when using TRITC filters (**Figure 3**). The increases in intracellular fluorescence when using FITC filters suggest at these excitation/emission wavelengths that quercetin stimulation causes autofluorescence likely due to intracellular permeation and/or alteration of fluorescent properties of intracellular components in these cell lines [42]. Notably, the lack of autofluorescence at the TRITC wavelengths used for Calbryte 590 calcium imaging demonstrates that this autofluorescence would not affect our calcium data, but caution may be warranted when imaging other fluorophores with FITC wavelengths in experiments with quercetin.

**Figure 3.**
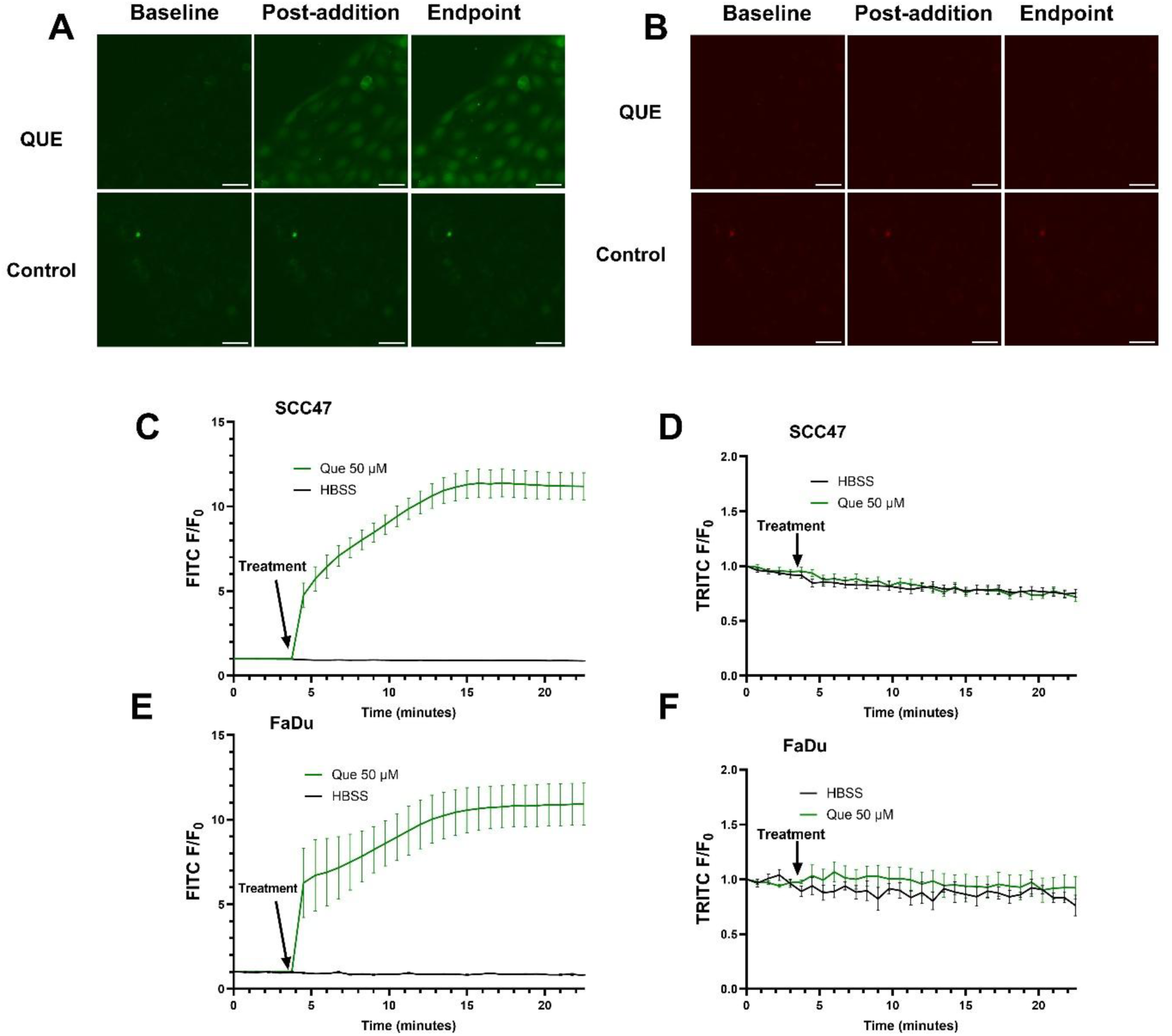
Autofluorescence detected in SCC47 and FaDu cells after addition of quercetin. Cells were washed and left in HBSS for imaging buffer, baseline images were acquired, and quercetin was added. FITC (474/27 nm_ex_, 525/45 nm_em_) and TRITC (554/23 nm_ex_, 609/54 nm_em_) filters were used to monitor autofluorescence signals. Cells were imaged at 20x at 45 second intervals. Increased signal may correspond to quercetin autofluorescence in cells. (**A**) Representative images in SCC47 cells using FITC and (**B**) TRITC filters. (**C**) Average traces in SCC47 using FITC and (**D**) TRITC filters. (**E**) Average traces in FaDu cells using FITC and (**F**) TRITC filters. Scale bars in representative images represent 50 µm. n = 4 technical replicates per condition with n = 3 biologic replicates in SCC47 and n = 1 biologic replicate in FaDu. Data points and error bars represent mean ± SEM.

### 3.3. Quercetin Decreases Cell Viability

To evaluate whether quercetin reduced cell viability in the studied HNSCC lines, crystal violet assays were performed on SCC47 cells and FaDu cells after exposure to varying concentrations of quercetin for 24 or 48 hours. Statistically significant decreases in viability were observed beginning at 100 µM in both cell lines at both timepoints (**Figure 4**). For SCC47, the IC_50_ values were 27.29 µM (95% CI: upper bound = 42.46 µM; lower bound could not be determined) and 37.43 µM (95% CI: 23.48-56.71 µM) at 24 and 48 hours, respectively. For FaDu cells, the IC_50_ values were 33.15 µM (95% CI: 24.27-40.84 µM) and 34.26 µM (95% CI: upper bound = 45.50 µM; lower bound could not be determined) at 24 and 48 hours, respectively. These findings suggest quercetin is cytotoxic to both HPV-positive and HPV-negative HNSCC cell lines at micromolar concentrations.

**Figure 4.**
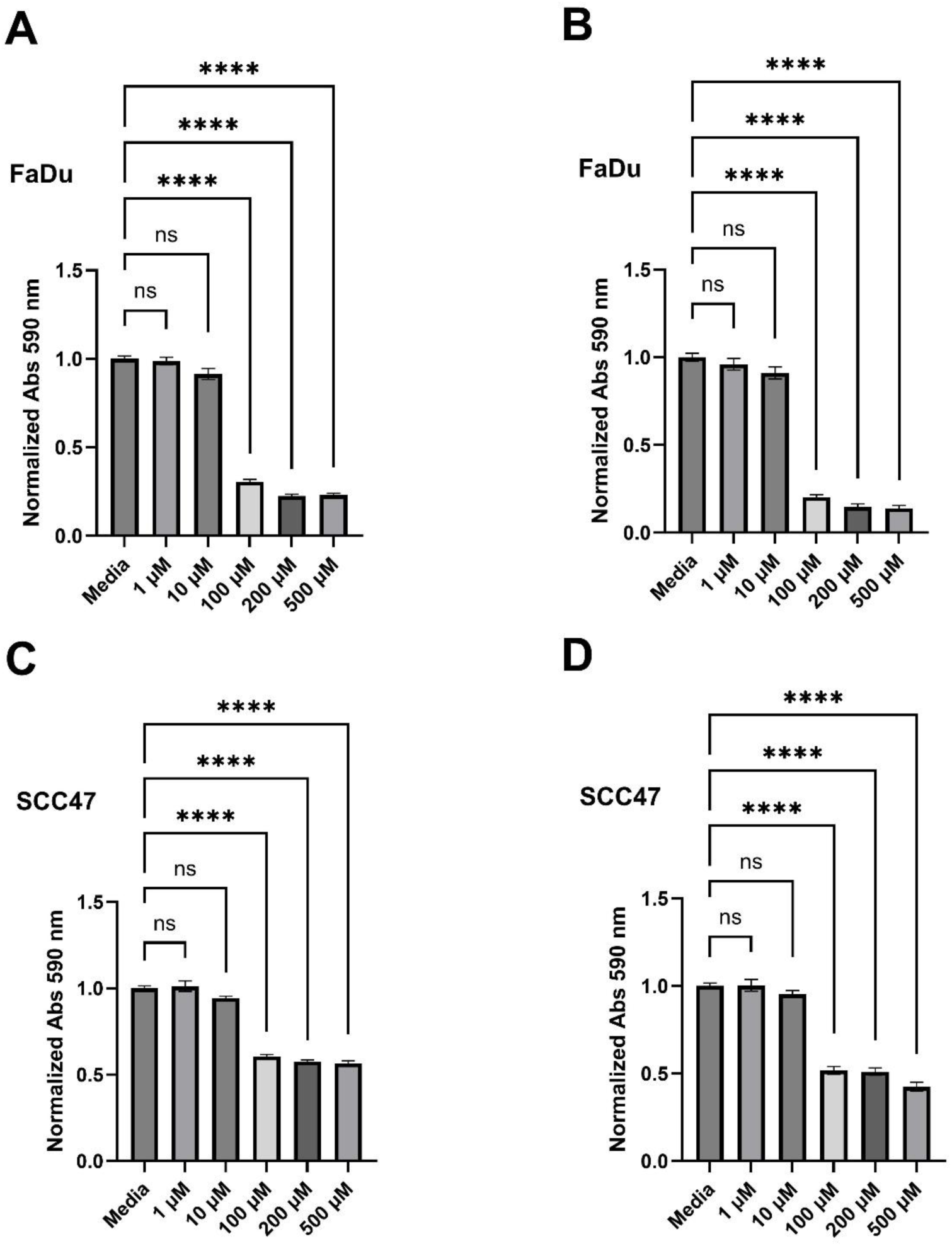
Quercetin decreases cell viability in HNSCC cell lines. Crystal violet assays were performed on SCC47 and FaDu cells treated with various concentrations of quercetin for either 24 or 48 hours. Media without quercetin was used as negative control in all experiments. (**A**) Cell viability after 24-hour and (**B**) 48-hour quercetin treatments in FaDu cells as assessed by crystal violet. (**C**) Cell viability after 24-hour and (**D**) 48-hour quercetin treatments in SCC47 cells assessed by crystal violet. Experiments were conducted with technical replicates with n = 3 and biologic replicates with n = at least 3. The statistical methods used were Brown-Forsythe and Welch ANOVA with Dunnet’s T3 multiple comparisons test (**A**) and ordinary one-way Anova with Tukey’s multiple comparison test (**B-D**). Data points and error bars represent mean ± SEM. ns, no statistical significance; ****P < 0.0001.

The effects of quercetin on viability were also assessed *ex vivo* using the MTS assay in patient-derived tumor slices. Following 24-hour exposure to 200 µM quercetin or control, the median of differences between control and treated samples was −0.31; however, Wilcoxon signed-rank test indicated that this difference was not significant, W = −11.00, P = 0.19. A before-after graph is shown in **Figure S2**. Given the limited sample size, additional studies with larger cohorts and extended exposure times are warranted to more definitively assess quercetin’s impact on tumor slice viability. Demographic and tumor information is shown in **Table S1**.

### 3.3 Quercetin Induces Mitochondrial Depolarization, Reduced by a T2R14 Antagonist

Mitochondrial membrane potential was monitored using TMRE. In SCC47 cells, TMRE minimum F/F_0_ signals were significantly reduced following quercetin treatment compared to control, with marked decreases observed within the first minute of quercetin exposure (**Figure 5**). These results indicate that quercetin induces rapid mitochondrial depolarization.

**Figure 5.**
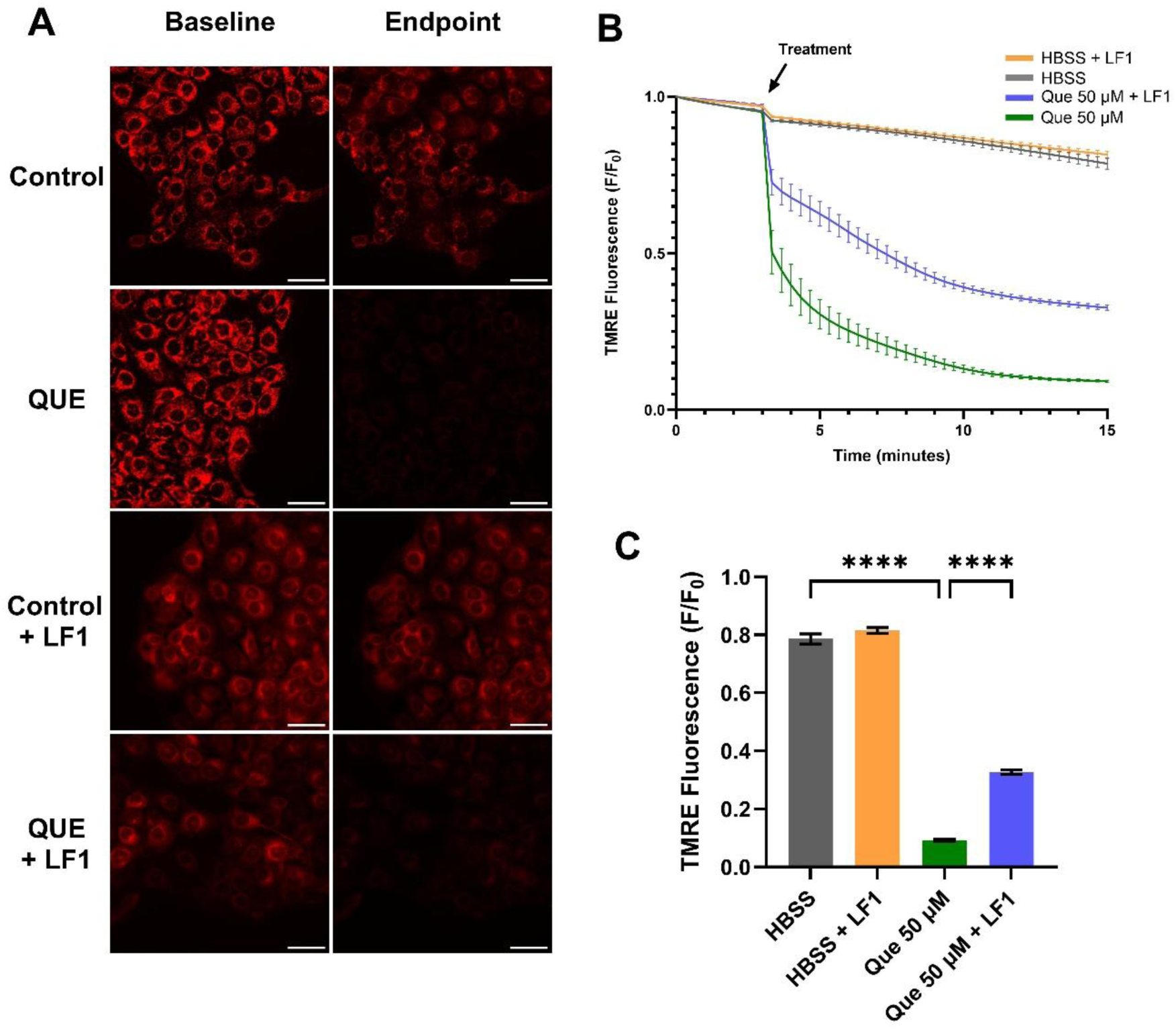
Mitochondrial depolarization in SCC47 cells exposed to quercetin with and without T2R antagonist LF1. Cells were incubated in TMRE and Hoechst dye with or without LF1 prior to imaging. Quercetin (with or without LF1) was added during imaging; HBSS or HBSS with LF1 served as negative controls. TMRE signal decreases when mitochondrial depolarization occurs. Cells were imaged at 20x using TRITC filters at 20 second intervals. (**A**) Representative images, (**B**) average traces, and (**C**) minimum TMRE F/F_0_ with and without LF1 in SCC47 cells are shown. Scale bars represent 50 µm. Experiments were conducted with technical replicates with n = at least 3 and biologic replicates = 3. Statistical method used was Brown-Forsythe and Welch ANOVA with Dunnet’s T3 multiple comparisons test. Data points and error bars represent mean ± SEM. ****P < 0.0001.

In the presence of the T2R14 antagonist LF1, mitochondrial depolarization was still observed, but minimum TMRE F/F_0_ signals after 15 minutes were significantly greater compared with quercetin alone (**Figure 5**). Similar findings were obtained in FaDu cells (**Figure S3**). Interestingly, in the presence of Gα_q/11_ inhibitor YM, quercetin did not cause significant differences in minimum TMRE F/F_0_ signals compared to control (**Figure S4**).

## 4. Discussion

In this study, we show that quercetin induces an intracellular Ca^2+^ response in HNSCC cells, which is significantly reduced in the presence of Gα_q/11_ inhibitor YM and T2R14 antagonist LF1. These findings support the role of quercetin as a T2R agonist, with T2R14 playing a central role in the observed Ca^2+^ responses. T2Rs interact with various G-proteins, with Ca^2+^ responses mostly dependent on Gβγ-mediated PLCβ activity [19, 25]. YM, a Gα_q/11_ inhibitor described as an “allosteric glue” that stabilizes the G-protein heterotrimer [43], reduced fluorescence signals, suggesting that quercetin-induced Ca^2+^ responses are likely mediated through Gα_q/11_ and associated Gβγ subunits. The reductions observed with LF1 further implicate T2R14 specifically, consistent with reports of LF1 as a selective T2R14 antagonist [26, 40].

Our findings expand upon prior reports that quercetin activates T2Rs. While initially identified as a T2R14 agonist, quercetin has since been shown to activate several additional receptors (e.g., T2R4, T2R7, T2R10, T2R31, T2R39, T2R40, T2R43, T2R46) in HEK293T cells, with the strongest Ca²⁺ response via T2R14 [33, 34]. In other cell types, such as human NCI-H716 intestinal L-cells and human choroid plexus papilloma cells, quercetin has been shown to enhance GLP-1 secretion via T2R38 [35] and trigger T2R14-mediated Ca²⁺ responses confirmed by knockdown [36]. Notably, in SCC47 and FaDu cells—which express multiple T2Rs [25, 26]— T2R14 antagonism alone was sufficient to significantly reduce quercetin-induced Ca²⁺ responses, suggesting that while quercetin may bind multiple receptors, its predominant effects in HNSCC cells are mediated through T2R14.

We also demonstrated autofluorescence in SCC47 and FaDu cells after addition of quercetin when using FITC filters, aligning with prior observations in HepG2 cells at similar excitation/emission ranges [42]. These results suggest intracellular accumulation of quercetin. Given that T2R14 has both extracellular and intracellular binding sites [44–46], these findings raise the possibility that quercetin’s effects may result from interactions at one or both sites.

Quercetin significantly decreased cell viability in SCC47 and FaDu cells, consistent with previous reports of quercetin’s cytotoxicity in various HNSCC lines [47]. In SCC47, absorbance values for conditions reducing viability were typically 50–60% of control, while in FaDu cells, viability reductions were greater at 20–30% of control. These differences between cell lines may reflect inherent variation in adherence and growth characteristics between the two lines or different susceptibility to quercetin.

*Ex vivo* studies using patient-derived tumor slices were more limited. Although the viability in some cases were decreased after treatment with quercetin, the results overall were not statistically significant with definitive conclusions potentially being limited by our sample size. Importantly, tumor slice models preserve elements of the tumor microenvironment—including immune cells and stromal cells [48]—which may influence therapeutic responses and the observed effects on viability when using the MTS assay. Future studies with larger cohorts are needed to determine whether quercetin decreases tumor slice viability and to evaluate potential effects on normal tissue.

Quercetin’s known ability to induce apoptosis may explain the reduced cell viability observed [49–53]. Prior studies in HNSCC and other cancers suggest contributions from mitochondrial depolarization and intrinsic apoptotic pathways [49, 51, 53–55]. Consistent with this, we observed quercetin-induced mitochondrial depolarization by TMRE imaging. LF1 partially mitigated this effect, whereas YM did not alter depolarization, suggesting that quercetin’s effects on mitochondrial membrane potential may involve mechanisms in addition to Gα_q/11_-mediated T2R signaling. Potential alternative mechanisms for quercetin-induced apoptosis include activation of mitochondrial large-conductance Ca^2+^-regulated potassium channels or interactions with the mitochondrial permeability transition pore [56, 57]. These findings do not definitively link T2R signaling to mitochondrial-driven decreases in viability during quercetin stimulation, and they underscore the complexity of quercetin’s mechanisms in HNSCC cells.

This study has limitations. Pharmacologic antagonists were used rather than T2R knockdown/knockout models, which would more definitively establish receptor specificity. Additional limitations include the small number of cell lines tested and small sample size for *ex vivo* tumor slices. Investigation in further cells lines, larger sample sizes for *ex vivo* models, and animal or patient-derived xenograft models are needed to better understand the potential T2R-mediated anti-cancer effects of quercetin in HNSCC tumors.

Overall, this work supports quercetin as a T2R agonist that induces intracellular Ca^2+^ responses in HNSCC cells, with T2R14 playing a key role. These findings contribute to the growing evidence for extra-oral T2Rs in cancer biology. Future studies using genetic knockdown or knockout approaches could more definitively define T2R14’s contribution. *Ex vivo* tumor slice models will be particularly valuable for validating mechanistic findings and assessing translational potential, especially given species differences between human and rodent T2Rs [34]. Finally, because many plant flavonoids and polyphenols are bitter compounds [58] with anticancer [3, 4] and anti-inflammatory properties [59], broader investigation of flavonoid T2R agonism in extra-oral tissues is warranted [60]. Beyond HNSCC, such studies may reveal novel roles for T2R signaling in normal physiology, disease pathophysiology, and therapeutic targeting.

## 5. Conclusions

This study demonstrates that quercetin induces intracellular Ca^2+^ responses in HNSCC cell lines, which are significantly reduced by the Gα_q/11_ inhibitor YM and the T2R14 antagonist LF1, supporting a key role for T2R14 in mediating these effects. Quercetin also decreases cell viability and promoted mitochondrial depolarization, adding to the evidence of its anticancer activity in HNSCC. Together, these findings identify quercetin as a functional T2R14 agonist in HNSCC cells and highlight the importance of extra-oral T2R signaling in tumor biology. While prior work has linked T2R activation to apoptosis, further studies are needed to clarify the mechanistic connection between quercetin-induced T2R signaling, mitochondrial dynamics, and cell death. These results suggest that targeting T2R14 signaling with quercetin or related compounds may represent a novel therapeutic strategy for HNSCC.

## Supplementary Materials

Additional Supplemental Figures can be found online in the Supplementary Material Tab. Figure S1: Live cell Ca^2+^ imaging in FaDu cells stimulated with quercetin in the presence and absence of Gα_q/11_ inhibitor; Figure S2: Effects of quercetin on cell viability of tumor slices; Table S1: Clinical data for HNSCC patients; Figure S3: Mitochondrial depolarization in FaDu cells exposed to quercetin with and without T2R Antagonist LF1; Figure S4: Mitochondrial depolarization assessed in SCC47 cells exposed to quercetin in the presence and absence of Gα_q/11_ inhibitor.

## Author Contributions

Conceptualization, Turner, G., Sywanycz, S., Hill, B., Wardlow, R., Lee, R., and Carey, R.; methodology, Turner, G., Sywanycz, S., Hill, B., Wardlow, R., Lee, R., and Carey, R.; formal analysis, Turner, G., Sywanycz, S., Hill, B., Wardlow, R., Lee, R., and Carey, R.; investigation, Turner, G., Sywanycz, S.; writing—original draft preparation, Turner, G.; writing—review and editing, Turner, G., Sywanycz, S., Hill, B., Wardlow, R., Lee, R., and Carey, R.; visualization, Turner, G.; supervision, Lee, R., and Carey, R.; All authors have read and agreed to the published version of the manuscript.

## Institutional Review Board Statement

The study was conducted in accordance with the Declaration of Helsinki, and approved by the Institutional Review Board of the University of Pennsylvania (protocol code 417200 approved 1/8/2025).

## Informed Consent Statement

Informed consent was obtained from all subjects involved in the study.

## Data Availability Statement

All data are contained within the manuscript, and other raw data are available upon request.

## Acknowledgments

We would like to acknowledge M. Victoria for her technical expertise and assistance.

## Abbreviations

The following abbreviations are used in this manuscript:

Ca^2+^: Calcium
DMEM: Dulbecco’s Modified Eagle Medium
DMSO: Dimethyl sulfoxide
FBS: Fetal Bovine Serum
GPCR: G-protein coupled receptor
HBSS: Hank’s Balanced Salt Solution
HNSCC: Head and neck squamous cell carcinoma
IP_3_: Inositol triphosphate
MTS: 3-(4,5-dimethylthiazol-2-yl)-5-(3-carboxymethoxyphenyl)-2-(4-sulfophenyl)-2H-tetrazolium
PLCβ: Phospholipase Cβ
T2R: Bitter taste receptor
TMRE: Tetramethylrhodamine ethyl ester
YM: YM-254980

