## Supplementary Materials for "Quercetin as a Bitter Taste Receptor Agonist with Anticancer Effects in Head and Neck Cancer Cells"

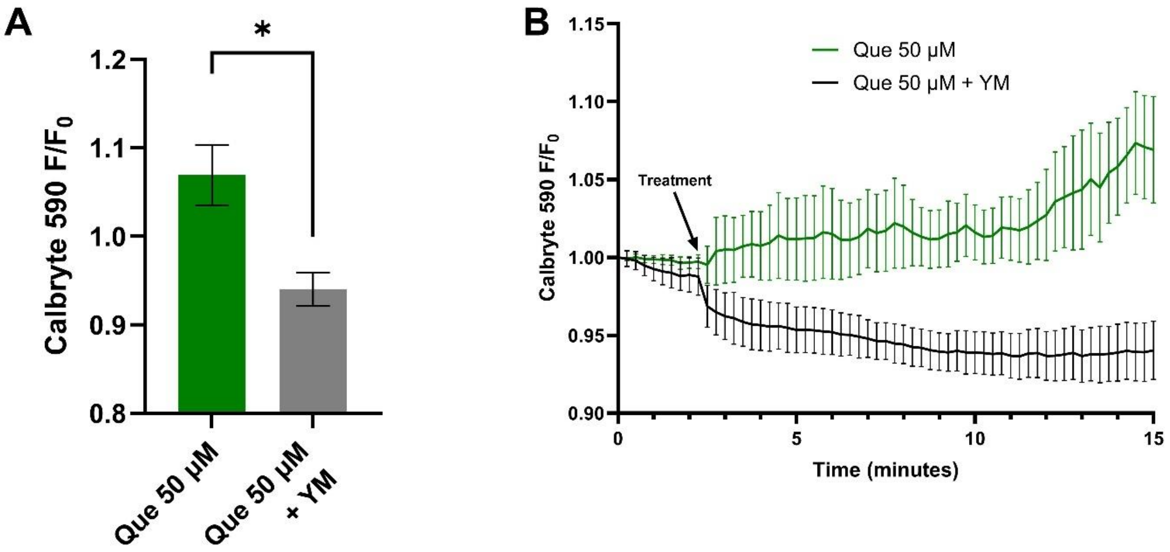


**Figure S1.** Live-cell $Ca^{2+}$ imaging in FaDu cells stimulated with quercetin in the presence and absence of $G\alpha_{q/11}$ inhibitor. FaDu cells were incubated in Calbryte 590 AM with and without YM prior to imaging. Quercetin with and without YM were added after baseline images were obtained; HBSS with or without YM served as the negative controls respectively. Cells were imaged at 20x using TRITC filters at 15 second intervals. Increased Calbryte 590 signals correspond to increased intracellular calcium. (**A**) Endpoint $F/F_0$ after approximately 11 minutes of stimulation with and without YM and (**B**) average traces in FaDu cells are shown. n = at least 3 technical replicates per condition in single experiment. The statistical method used was unpaired t test. Data points and error bars represent mean ± SEM. *P < 0.05.

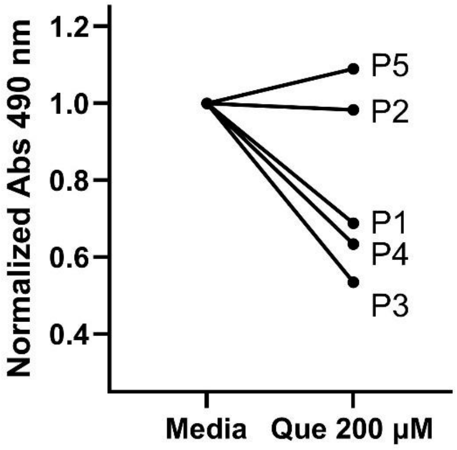


**Figure S2**. Effects of quercetin on cell viability of tumor slices. Tumor slices derived from patients were treated with quercetin or media control for 24 hours. Cell viability was assessed with the MTS assay and measuring absorbance at 490 nm, with lower values corresponding with less viability. Shown is a before-after graph. Each data point represents the averaged absorbance values normalized to respective controls per patient. Three technical replicates were performed per condition with n = 5 patients ("P1" corresponding to patient 1 in Table S1, "P2" to patient 2, etc.).

**Table S1**. Clinical data for HNSCC patients.

| Patient | Age | Sex | Race/ethnicity | Location of Primary Tumor | p16 status | Prior H&N Treatment History |
|---|---|---|---|---|---|---|
| 1 | 66 | Male | Unknown/non-Hispanic | Tonsil (palatine) | + | none |
| 2 | 55 | Male | White/non-Hispanic | Oral/gingivobuccal | - | none |
| 3 | 51 | Male | White/non-Hispanic | Lateral tongue | Not reported | none |
| 4 | 77 | Male | White/non-Hispanic | Tonsil (palatine) | + | Radiation, (this new primary lesion was outside of radiation field) |
| 5 | 67 | Male | White/non-Hispanic | Oropharynx (base of tongue) | + | Definitive chemoradiation approx. 3 years prior |

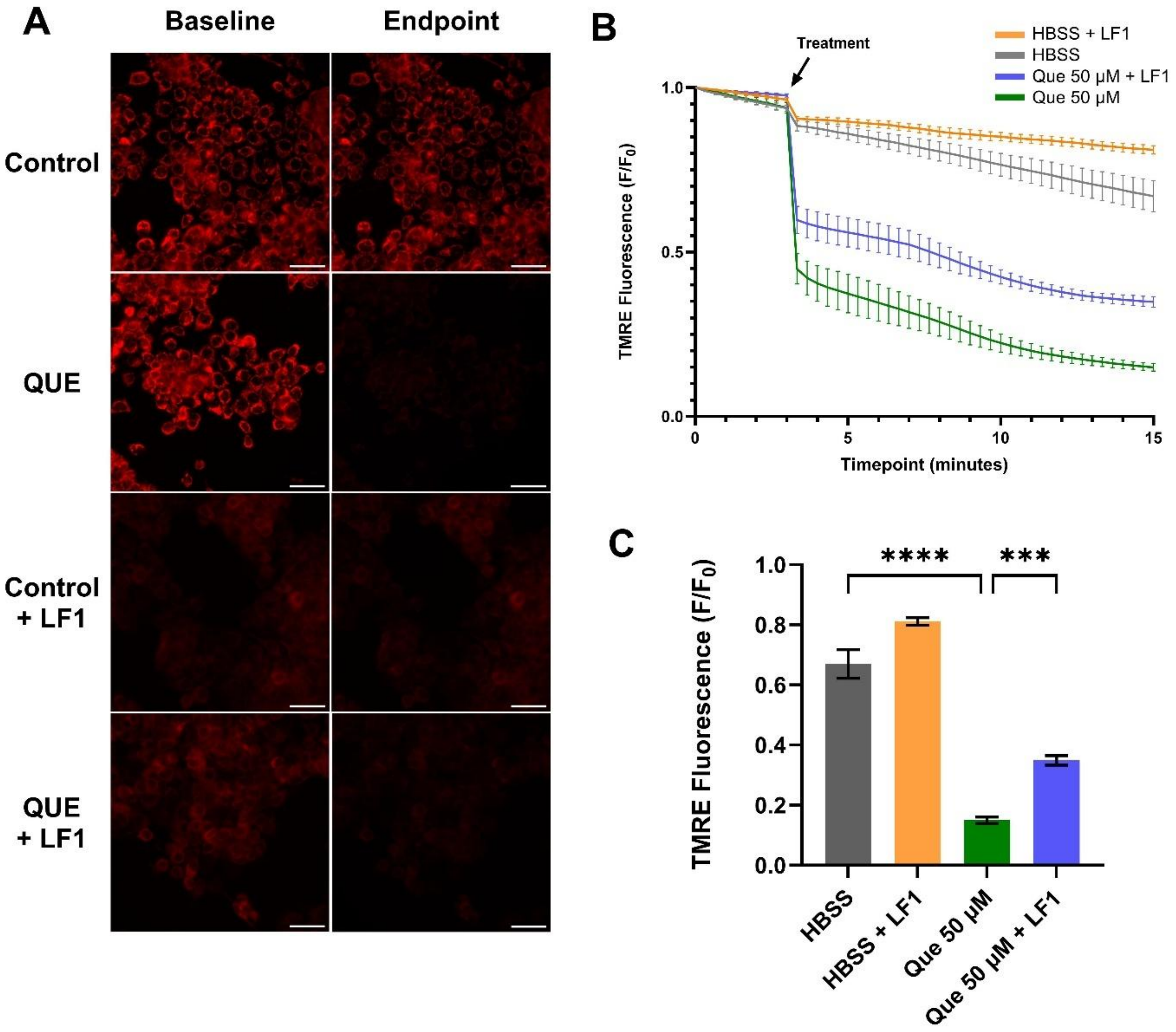


**Figure S3.** Mitochondrial depolarization in FaDu cells exposed to quercetin with and without T2R Antagonist LF1. Cells were incubated in TMRE and Hoechst dye with or without LF1 prior to imaging; quercetin with or without LF1 was added during imaging and HBSS or HBSS with LF1 was used for negative controls. TMRE signal decreases when mitochondrial depolarization occurs. Cells were imaged at 20x using TRITC filters at 20 second intervals. (**A**) Representative images, (**B**) average traces, and (**C**) minimum TMRE $F/F_0$ with and without LF1 in FaDu cells are shown. Scale bars in representative images represent 50 µm. Single experiment was conducted with technical replicates n = 4. Statistical method used was ordinary one-way Anova with Tukey's multiple comparison test. Data points and error bars represent mean ± SEM. $^{***}P < 0.001$; $^{****}P < 0.0001$.

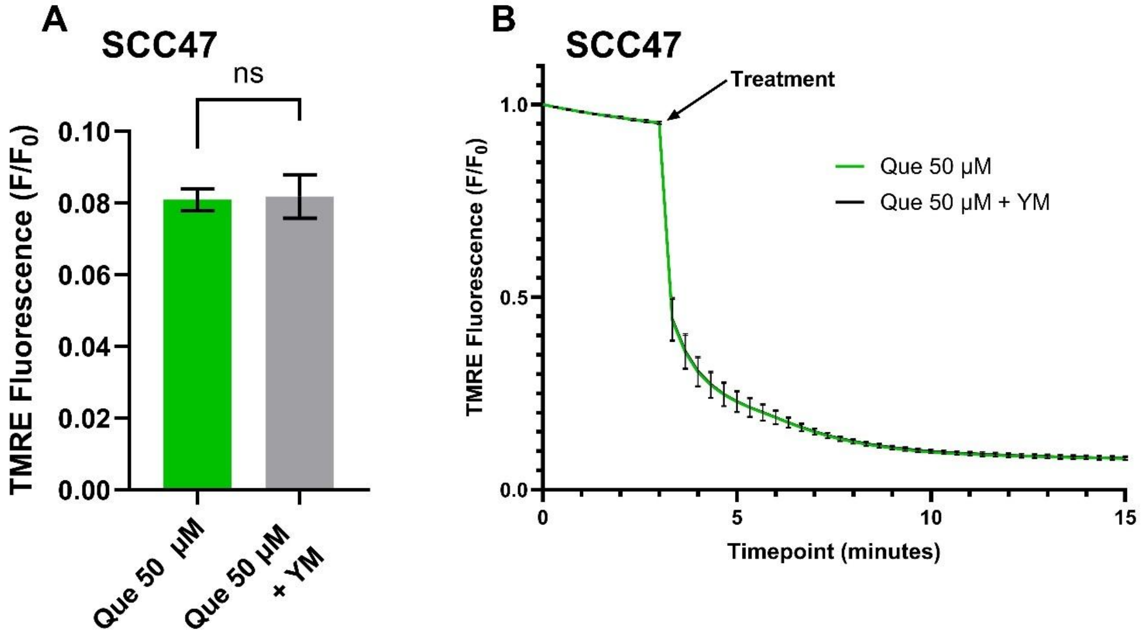


**Figure S4.** Mitochondrial depolarization assessed in SCC47 cells exposed to quercetin in the absence and presence of G$\alpha_{q/11}$ inhibitor. Cells were incubated in TMRE and Hoechst dye with or without YM prior to imaging. Quercetin with or without YM was added during imaging and HBSS or HBSS with YM was used for negative controls. TMRE signal decreases when mitochondrial depolarization occurs. Cells were imaged at 20x with 50 ms exposure and 12.5% LED intensity using TRITC filters at 20 second intervals. (**A**) Minimum TMRE $F/F_0$ and (**B**) average traces in SCC47 cells exposed to quercetin with and without YM. n = 4 technical replicates in a single experiment. Statistical method used was unpaired t test. ns, no statistical significance.
